# AAV-mediated gene transfer of WDR45 corrects neurologic deficits in the mouse model of beta-propeller protein-associated neurodegeneration

**DOI:** 10.1101/2024.06.18.599588

**Authors:** Maria Carla Carisi, Claire Shamber, Martha Bishop, Madison Sangster, Uma Chandrachud, Brandon Meyerink, Louis Jean Pilaz, Yulia Grishchuk

**Affiliations:** Center for Genomic Medicine and Department of Neurology, Massachusetts General Hospital Research Institute and Harvard Medical School, Boston, MA, USA; Pediatrics and Rare Diseases Group, Sanford Research Institute, University of South Dakota, Sioux Falls, USA

**Keywords:** WIPI4, WDR45, autophagy, adeno-associated virus, autophagy, gene transfer, gene addition

## Abstract

Beta-propeller protein-associated neurodegeneration (BPAN) is an ultra-rare, X-linked dominant, neurodegenerative disease caused by loss-of-function mutations in the *WDR45* gene. It manifests in neurodevelopmental delay and seizures followed by secondary neurologic decline with dystonia/parkinsonism and dementia in adolescence and early adulthood and is characterized by progressive accumulation of iron in the basal ganglia. *WDR45* encodes β-propeller-shaped scaffold protein, or WIPI4, which plays an important role in autophagosome formation. While the mechanisms of how WIPI4 loss of function results in neurologic decline and brain pathology have not yet been established, findings of lower autophagic activity provide a direct link between impaired autophagy and neurologic disease in BPAN. Here we performed phenotypical characterization of a novel mouse model of BPAN, WDR45_ex9+1g>a mouse. We identified hyperactive behavior and reduction of autophagy markers in brain tissue in *WDR45*_ex9+1g>a hemizygous males as early as at 2 months of age. Given the early onset and spectrum of neurologic symptoms such as hyper-arousal and attention deficits in human patients, this model presents a disease-relevant phenotype and can be used in preclinical studies. We used this mouse model for a proof-of-concept study to evaluate whether AAV-mediated CNS-targeted gene transfer of *WDR45* can provide therapeutic benefit and be considered a therapeutic paradigm for BPAN. We observed successful expression of human *WDR45* transcripts and WIPI4 protein in the brain tissue, rescue of hyperactive behavior, and correction of autophagy markers in the brain tissue. This data demonstrates that *WDR45* gene transfer can be a promising therapeutic strategy for BPAN.

## Introduction

Beta-propeller protein-associated neurodegeneration (BPAN, OMIM:300894), also known as a neurodegenerative disease with brain iron accumulation type 5 (NBIA5) and static encephalopathy of childhood with neurodegeneration (SENDA), is an ultra-rare, X-linked dominant, neurodegenerative disease caused by pathogenic mutations in the *WDR45* gene (1, 2). The clinical manifestations of BPAN are highly heterogeneous between individuals, and the disease is significantly more prevalent in females, with only a few male patients identified to date, likely due to higher severity and embryonic lethality in the hemizygous state (3, 4). Individuals with BPAN are phenotypically diverse and may present with a global severe neurologic disability, relatively preserved gross motor function but severely impaired language development, or have an attenuated phenotype, demonstrating measurable language skills and spared neurologic function (5). During early childhood, other symptoms include seizures and cognitive impairment such as repetitive movements, bruxism, limited attention span, hyper-arousal, restlessness, and abnormal behaviors. Later in life, starting in adolescence or young adulthood, the onset of progressive movement disorder in the form of dystonia and Parkinsonism, and dementia has been documented (5, 6, 7, 8). The main brain MRI findings are hypomyelination and thinning of the corpus callosum, which can be seen in some but not all patients, and a more common progressive deposition of iron in the basal ganglia (9). Brain pathology evaluation is limited to two published cases and includes neuronal loss, predominantly cerebral atrophy, gliosis, axonal swellings, Alzheimer ’s-like Tau-tangles, and iron deposits in the basal ganglia (6, 10). The complexity of the clinical manifestations and the rarity of the disease contribute to the difficulty of obtaining a diagnosis. BPAN can be misdiagnosed as autism, epilepsy, or other forms of NBIA (9). Currently, there are no approved or experimental therapies for BPAN, and treatment options are limited to only symptom management.

WDR45 gene encodes the WD repeat domain phosphoinositide-interacting protein 4 (WIPI4), one of four proteins of the PROPPIN (β-propellers that bind polyphosphoinositides), also known as WIPI (WD repeat domain, phosphoinositide interacting) family. WIPI proteins function as membrane remodeling proteins with a central role in autophagosome formation in autophagy, a metabolic process of recycling intracellular materials via lysosomal degradation necessary for cell survival. The initiating step of the autophagic process is the formation of the phosphoinositide-3 kinase class III (PI3KC3) complex I composed of VPS34, VPS15, Beclin 1, and ATG14L, and the consequent production of phosphatidylinositol 3-phosphate (PtdIns3P, or PI3P) (11, 12). PI3P has a dual function in autophagophore formation. First, it controls the signaling of autophagy initiation, and second, it is used for building phagophore membrane fragments in the ER subdomains called omegasomes (13, 14, 15, 16, 17). The four mammalian WIPI proteins, WIPI1-4, act as PI3P effectors that decode PI3P signals and consequently enable autophagophore formation, working as scaffolds and attracting their protein partners to the forming autophagosomal membrane. WIPI4 forms a complex with ATG2 that recruits phospholipids from the endoplasmic reticulum to expand the autophagic membrane (18, 19, 20). Mutations of the ATG2A binding sites in WIPI4 disrupt autophagic flux (21) and have been documented in BPAN patients, where the severity of the disease significantly correlates with the inhibitory effect of these mutations (22). Over 90 *WDR45* variants associated with BPAN have been documented, with no specific protein domain linked to the disease, most resulting in complete protein loss (3). While the precise pathophysiological mechanisms of how loss of *WDR45* results in neurologic decline and brain pathology, including brain iron accumulation, have not yet been established, findings of lower autophagic activity and accumulating autophagic vacuoles provide a direct link between impaired autophagy and neurologic disease (2).

Several mouse models of *WDR45* deficiency have been created to date, including constitutive (23), (24), (25) and conditional, pan-neuronal (26), and dopaminergic neuron-specific *WDR45* knock-out model (27). While these models reproduce many molecular, cognitive, and neurologic manifestations of BPAN and the progressive course of the disease, motor and congnitive deficits develop in these mouse models late in life, and do not replicate neurodevelopmental aspects of the disease seen in human patients. This late presentation of disease phenotype in these BPAN mouse models can present challenges for their use in translational nonclinical studies since most human patients have neurodevelopmental phenotypes, and early interventions would be required for the optimal therapeutic strategy in this disease. Therefore, in this study, we have characterized another mouse model of BPAN, a Wdr45_ex9+1g>a mouse, available via the Mutant Mouse Resource & Research Centers (MMRRC) repository. We identified hyperactive behavior and reduction of autophagy markers in brain tissue in Wdr45_ex9+1g>a hemizygous males as early as at 2 months of age. Using this mouse model, we have performed proof-of-concept studies of AAV-mediated *WDR45* gene transfer to rescue neurologic dysfunction and autophagy deficits in BPAN. Our data show that AAV-mediated expression of human *WDR45* was efficacious to rescue disease phenotype in BPAN mice, proving that this approach can be a promising therapeutic strategy for BPAN and opening the door to translational development.

## Materials and Methods

### Animals

The WDR45_ex9+1g>a mice on the C57BL/6N background were obtained from the Mutant Mouse Resource and Research Center (MMRRC) national repository (RRID: 067175-UCD) and housed according to National Institutes of Health (NIH) guidelines for animal care under the approval of the Animal Care and Use Committee (IACUC). All procedures performed were approved by the Massachusetts General Hospital Subcommittee on Research Animal Care. Wild-type littermates were used as controls.

Mice were genotyped using the Transnetyx genotyping services (Cordova, TN) using qRT-PCR (https://www.transnetyx.com/). The recessive RD8 mutation of the Crb1 gene has been previously identified in the C57BL/6N strain (28), causing retinal degeneration and eye depigmentation. Crb1 RD8 allele was present in the breeders obtained from MMRRC and was removed from our experimental cohorts by crossing C57BL/6N WDR45_ex9+1g> hemizygous males (Wdr45^y/ex9+1g>a^) with the wild type C57BL/6J (Rd8 negative) mice for at least 5 generations. We have observed that Wdr45^y/ex9+1g>a^ Crb1 ^Rd8/Rd8^ males obtained from the original breeders for phenotype characterization developed excessive body mass after 6 months of age, and all Crb1 ^Rd8/Rd8^ males have been excluded from the study.

### Constructs and AAV preparation

A self-complimentary pAAVsc-JeT-WDR45 expression plasmid was used to obtain a scAAV9-JeT-WDR45, and a single-stranded pAAVss-JeT-WDR45 expression plasmid was used to produce AAV-PHP.eB- JeT- *WDR45* viral particles for this study. The expression vector backbones were obtained from Dr Luk Vandenberghe’s lab at Massachusetts Eye and Ear Infirmary. A human *WDR45* cDNA, polyA (29), and minimal synthetic JeT (30) promoter were synthesized and subcloned to construct the expression cassette. The obtained rAAV cis constructs were verified by whole plasmid sequencing via NanoPore technology. Viral production was performed under research-grade conditions at Signagen (Frederick, MD) (https://signagen.com/). To package the rAAVs, the HEK293T cells were seeded in Cell Stacks at 3x10E+5 cells/mL and transiently transfected with a helper-free rAAV packaging system. Specifically, the obtained rAAV cis construct, the corresponding capsid plasmid, and the helper plasmid were co-transfected to HEK293 cell via PolyJet™ In Vitro DNA Transfection Reagent (SignaGen, Frederick, MD). Three days after transfection, cells were detached and harvested using PBS buffer. Harvested cells were lysed by three freeze/thaw cycles before an overnight Benzonase treatment at 37 °C to digest residual DNA followed by high salt treatment for 1h at 37 °C. The crude lysate was further clarified by low-speed centrifugation. Full capsid AAVs were purified by ultracentrifugation in a cesium chloride (CsCl) gradient. A diafiltration step was then performed to concentrate the purified AAVs and reformulate them into PBS + 0.005% poloxamer 188 (Corning, <u>Manassas</u>, VA) using Amicon Ultra-4 filter units (Millipore, <u>Burlington, MA</u>) with a 100 kDa MWCO membrane. Finally, a sterile filtration through a 0.221μm membrane filter was performed. Vector genome titer was determined by qPCR using an ITR-specific primer-probe set.

The final purified products were characterized with assays to measure identity, potency, and purity. scAAV9-MCOLN1 vector was used from the previously described batch (31).

### ICV injections

The viral particle stock was diluted in sterile saline (0.9% NaCl) (Hospira, Lake Forest, IL) with 0.01% Fast Green (Sigma, St. Louis, MI). A 10-µl Hamilton syringe (Hamilton Company, Reno, NV) with a 30G needle was used to inject each pup with 5 µl of either saline, scAAV9-JeT-WDR45 at a dose of 3e10 vg/mL, or scAAV9-JeT-MCOLN1 at a dose of 2e10 vg/mouse. Litters were randomly assigned to either “saline”, “WDR45-treated,” or “MCOLN1-treated” groups. Injections were performed as previously described (31). Briefly, pups were anesthetized on ice for 5-7 minutes, then placed on an LED lamp to make skull anatomy visible, and injections were performed about 0.25 mm lateral to sagittal suture and 0.50–0.75 mm rostral to neonatal coronary suture (0.25 mm lateral to confluence of sinuses). After injection, pups were placed under a warming lamp and monitored for recovery, then moved back with the mother when movement and responsiveness were fully restored.

### IV injections

Mice over 16 g of body weight were used for tail-vein injections. Cages with male mice were randomly assigned to either “saline” or “treated” groups. Mouse tail veins were dilated by exposure to warm light for a few minutes, and animals were confined in a plexiglass restrainer. Injections were performed using 30G-needle insulin syringes (Cat. No 328466, BD, San Diego, CA) with 90 or 100 µL of either saline or AAV-PHP.eB-Jet-WDR45 at a dose of 7.2×10e11 vg/mouse. After injection, mice were returned to their home cages and monitored daily.

### In-vivo mouse testing

All procedures were performed by the investigator blind to the genotype and treatment group information. Open field testing was performed on naive male mice at 2, 6, or 11 months of age under regular light conditions, as previously described (31). Briefly, each mouse was placed in the center of a 27×27 cm^2^ plexiglass arena and recorded by the Activity Monitor program (Med Associates Fairfax, VT). The horizontal and vertical activities were analyzed during the first 15 min in the central (8×8 cm^2^) versus peripheral (residual) zone of the arena. Motor coordination and balance were tested on an accelerating rotarod for two days (Med Associates, VT). Latency to fall from the rod (accelerating speed from 4 to 40 rpm over 5 min) was recorded in 3 trials on the second day. Neurological exams consisted of 13 steps: assessment of general health, righting reflex, crossed extensor reflex, forelimb and hindlimb placing responses, grasp reflex, postural reflex, rooting reflex, placing response, vibrissae orientation, visual placing response, eyeblink response, negative geotaxis, and acoustic startle response.

### Tissue collection and processing

Mice were sacrificed using a carbon dioxide chamber. Immediately after euthanasia, mice were transcardially perfused with ice-cold phosphate-buffered saline (PBS). After bisecting the brain across the midline, the cortex was isolated from half and snap-frozen over dry ice before placing it for storage at −801C.

### Protein extraction and SDS-PAGE Western blotting

Mouse cortical tissue was manually disrupted using homogenization pesters (Fisher Scientific, Hampton, NH, 13236679) in the lysis buffer [20 mmol HEPES pH 7.4 (Sigma, St. Louis, MI H0763), 10 mmol NaCl (Sigma, St. Louis, MI S7653), 3 mmol MgCl2·6H2O (Sigma, St. Louis, MI M2670), 2.5 mmol EGTA (Sigma, St. Louis, MI E3889), 50 mmol NaF (Sigma, St. Louis, MI S7920), 100mM DTT (Bio-Rad, Hercules, CA, #161-0611), 1% Igepal ca-360 (Sigma, St. Louis, MI, I-8896), and a protease inhibitor cocktail (Calbiochem, San Diego, CA, #539134)]. Lysates were incubated on ice for 45 minutes, vortexed 3 times with 15 min intervals, and centrifuged at 12000 rpm for 5 minutes at 4C to remove debris. Protein concentration was measured using Pierce™ BCA Protein Assay Kit (Thermo Fisher, Waltham, MA, 23227). 30µg of protein homogenates were separated by SDS-PAGE on a 4-12% gradient Bis-Tris gel (Bio-Rad, Hercules, CA, Criterion XT 3450125) in XT MES buffer (Bio-Rad, Hercules, CA, 1610796) and transferred for 7 minutes on a nitrocellulose membrane (Bio-Rad, Hercules, CA, 1620233) in XT transfer buffer (Bio-Rad, Hercules, CA, 10026938). Membranes were incubated overnight with the following primary antibodies: αWIPI2 1:3000 (mouse, Bio-Rad, Hercules, CA, MCA5780GA), αWIPI4/WDR45 1:2000 (rabbit, Proteintech, Rosemont, IL, 19194-1-AP), αLC3B 1:500 (rabbit, Sigma, St. Louis, MI, L8919), αActin 1:1000 (rabbit, Sigma, St. Louis, MI, a2066), αActin 1:5000 (mouse, Sigma, St. Louis, MI, a5441). The following secondary antibodies were applied: αRabbit IRDye 800 1:10,000 (goat, LI-COR, Lincoln, NE, 926-32211), αMouse IRDye 800 1:10,000 (goat, LI-COR, Lincoln, NE, 926-32210), αRabbit IRDye 680 1:20,000 (goat, LI-COR, Lincoln, NE, 926-68071), αMouse IRDye 680 1:20,000 (goat, LI-COR, Lincoln, NE, 926-68070).

Proteins were visualized using the Odyssey Infrared Imaging System (LI-COR, Lincoln, NE). Image Studio v5.2 software (LI-COR, Lincoln, NE) was used for densitometric analysis, and OD values were normalized to Actin.

### RNA extraction and qPCR analysis

Mouse brain tissues were disrupted and homogenized using QIAzol lysis reagent (Qiagen, Hilden, Germany, 79306), and the Tissue Lyser instrument (Qiagen, Hilden, Germany); snap-frozen tissue was processed by adding 500 µl of QIAzol reagent in the presence of one 5-mm stainless steel bead (Qiagen, Hilden, Germany). Total RNA isolation from homogenized tissues was performed using a Qiagen RNeasy kit (Qiagen, Hilden, Germany), and genomic DNA was eliminated by DNase (Qiagen, Hilden, Germany) digestion on columns following procedures indicated by the provider. cDNA was generated using a High-Capacity cDNA Reverse Transcription kit (Applied Biosystems, Foster City, CA, 4368814) according to manual instructions. cDNA produced from 500 ng of starting RNA was diluted, and 40 ng was used to perform qPCR using Light Cycler 480 Probes Master mix (Roche Diagnostics, Mannheim, Germany, 04707494001). The real-time PCR reaction was run on Light Cycler 480 (Roche Diagnostics, Mannheim, Germany) using TaqMan premade gene expression assays (Applied Biosystems, Foster City, CA) and using the following probe sets: Mus musculus GFAP (FAM)-Mm01253033_m1, CD68 (FAM)- Mm03047343_m1, and GAPDH (FAM)-Mm99999915_g1. Quantitative qRT-PCR to measure *WDR45* expression was performed using *WDR45* (FAM)-Hs01079049_g1 probe set (Applied Biosystems, Foster City, CA). To obtain a standard curve, the scAAV-JeT-WDR45 plasmid was linearized with HindIII (New England Biolabs, Ipswich, MA, USA) as directed by the manufacturer. The DNA concentration was determined after digestion, and the number of copies of DNA/ul was calculated assuming a molar mass of 650 g/mol per base pair and a fragment length of 4500 nt. A standard curve with serial dilutions of the linearized plasmid ranging from 4 x 10^2^ to 4 x 10^7^ copies per reaction was used to determine the absolute number of cDNA transcripts. To quantify MCOLN1 (TRPML1) transcripts, three TaqMan probes were designed: forward 5’-GGTCGCGGTTCTTGTTTGT-3’, reverse 5’- GAAGCCGCTCGGTCTCT-3’, probe 5’- CCCTGTGATCGTCACTTGACAGTGT-3’. The scAAV-JeT-MCOLN1 plasmid was linearized with HindIII (New England Biolabs, Ipswich, MA, USA) to obtain a standard curve. The DNA concentration was determined after digestion, and the number of copies of DNA/ul was calculated assuming a molar mass of 650 g/mol per base pair and a fragment length of 5158 nt. A standard curve with serial dilutions of the linearized plasmid ranging from 4 x 10^2^ to 4 x 10^7^ copies per reaction was used to determine the absolute number of cDNA transcripts.

### Vector genome quantification

Vector genome copy number was quantified by qPCR analysis of total DNA extracted from mouse tissues using Qiagen DNeasy Blood and Tissue kit and Tissue Lyser II (Qiagen, Hilden, Germany). Genomic RNA was eliminated by RNase digestion (Qiagen, Hilden, Germany); 40 ng per sample at 10 ng/µl for cortical samples were used to perform qPCR using Light Cycler 480 Probes Master mix (Roche Diagnostics, Mannheim, Germany) on Light Cycler 480 (Roche Diagnostics, Mannheim, Germany) using a custom probe/primer set (ITRJET(FAM)-APMF3G9) created via Custom TaqMan® Gene Expression Assays Design Tool (Applied Biosystems, Foster City, CA) and designed to bind to a fragment of the expression vector including 3’-ITR and JeT promoter sequence. The standard curve with serial dilutions of the pAAVsc-JeT- *WDR45* plasmid ranging from 1e7 ng/µl to 1e2 ng/µl was used to determine the absolute number of copies of the viral genome/µg of total DNA, and the values were converted to vector copies per mouse genome assuming an average mass of 5.4 pg per a diploid mouse genome.

### Statistical analysis

All data are expressed as individual values, mean ± SEM, or mean ± SEM. Analyses were performed in Prism version 10 (GraphPad, La Jolla, CA), using the ROUT (Q=1) method for identifying outliers, unpaired T-test, One-way or Two-way ANOVA with correction tests for multiple comparisons as detailed for specific experiments in figure legends. P values were indicated as follows throughout the manuscript: n.s. = p > 0.05; * p =< 0.05; ** p =< 0.01, *** p = < 0.001,**** p =< 0.0001.

## Results

*Wdr45*^ex9+1g>a^ mouse model of BPAN shows disease-relevant phenotype.

To characterize neurologic deficits in Wdr45_ex9+1g>a mouse model of BPAN from MMRRC (RRID:MMRRC_067175-UCD), we employed the open-field test that provides quantitative measurements of spontaneous and exploratory activity and locomotion, and the rotarod test that measures motor function, balance, and coordination (Fig.1A, cohort 1). We found that BPAN (Wdr45 ^ex9+1g>a/y^, later referred to as Hemi) male mice display hyperactivity behavior in the open field test at two months of age, shown as higher values of the total distance traveled, ambulatory time (time spent moving), ambulatory episodes (number of times movement was initiated) and ambulatory counts (number of movement events) (Fig. 1B). Since the open field test measures spontaneous activity in mice, testing of naïve mice is often required to observe phenotypical differences. Therefore, to test whether the hyperactivity behavior we have observed in young adult mice at 2 months would persist in the course of the disease, we have generated an additional cohort of mice that were subjected to an open field test only at 11 months of age (Fig 1A, cohort 2). Aged BPAN mice also demonstrated a significant increase in track length, ambulatory time, and ambulatory episodes, indicative of hyperactivity (Fig. 1B). Next, we used a monthly rotarod test in Cohort 1 animals to continuously assess motor function, balance, and coordination in the course of disease. BPAN Hemi males did not show a significant difference from their healthy controls (Fig. 1C). In parallel with the rotarod testing, we have also employed monthly weighing (Fig. 1D) and evaluation of basic neurologic responses (i.e. neuroexam; not shown) in BPAN mice and found no significant changes.

**Figure 1.**
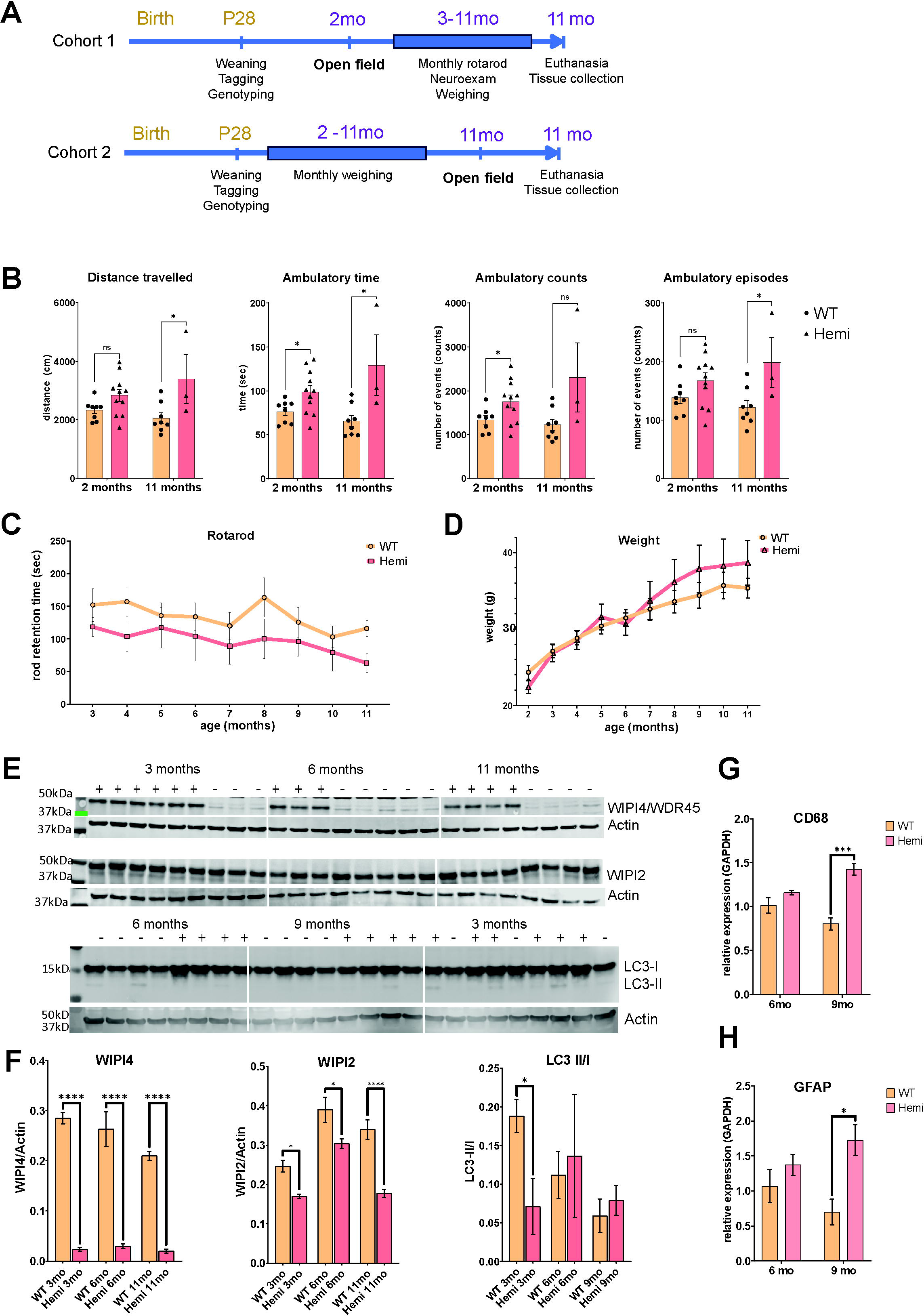
Characterization of disease phenotype in Wdr45^ex9+1g>a/y^ male mice. A. Schematic representation of the experimental design. B. Wdr45^ex9+1g>a/y^ (Hemi) mice show hyperactive behavior in the open field test. Data are shown as individual values and average ± SEM, * p<0.05; **p<0.01; Unpaired T-test; n= 8 (WT, 2mo), n=11 (Hemi, 2 mo), n= 8 (WT, 11 mo), n= 3 (Hemi, 11 mo). C. Monthly rotarod testing shows no significant changes between Hemi and WT male mice from 3 to 11 months of age. Data are shown as average values ± SEM; n (WT) = 6 (3-10 mo) and 14 (11 mo), n (Hemi) = 6 (3-10 mo) and 9 (11 mo). Grouped statistical analysis using a mixed-effects model with the Geisser-Greenhouse test (GraphPad Prizm): p (age)= 0.0043; p (genotype) = 0.087. Sidak’s multiple comparisons test showed no significant differences between WT and Hemi at any time point. D. No significant changes in body mass have been observed between Hemi and WT mice from 2 to 11 months of age. Data are shown as average values ± SEM; n (WT) = 3 (2 mo), 10 (3-10 mo) and 14 (11 mo), n (Hemi) = 5 (2mo), 7 (3-10 mo) and 9 (11 mo). Grouped statistical analysis using a mixed-effects model with the Geisser-Greenhouse test (GraphPad Prizm): p (age) <0.0001; p (genotype) = 0.94. Sidak’s multiple comparisons test showed no significant differences between WT and Hemi at any time point. Western blot (E) and quantification (F) of Wipi4, Wipi2 and Lc3 proteins in cerebral cortex of Hemi (-) and WT (+) mice at 3, 6 and 9 or 11 months of age. Data shown as average ± SEM, * p<0.05; ****p<0.0001 by Ordinary one-way ANOVA and Sidak’s test for multiple comparisons; n= 6 (WT, 3 mo), n=3 (Hemi, 3 mo), n= 3 (WT, 6 mo), n= 5 (Hemi, 6 mo), n= 4 (WT, 11 mo), n= 4 (Hemi, 11 mo). qRT-PCR analysis of the activated microglia/macrophage marker CD68 (G) astrocyte marker GFAP (H) showed an increase in the Hemi cortex at 9 months. Data are shown as average ± SEM, * p<0.05; ***p<0.001; two-way ANOVA (p (age) = 0.32; p (genotype)<0.0001) and Tukey multiple comparisons test; n= 4 per genotype and age group.

Western blot (Fig. 1E and F) on brain protein homogenates showed reduced Wipi4 expression in Hemi mice. Analysis of the autophagosome formation marker LC3 showed a decreased ratio of LC3-II/LC3-I at 3 months, suggesting inhibition of autophagy. We also found a significant reduction in the levels of Wipi2 in Hemi brain tissue at 3, 6, and 11 months of age. Wipi2 is another WIPI protein family member interacting with Wipi4 during autophagosome formation. Other markers of autophagy activation, such as beclin 1 or p62/SQSTM, did not show any difference between genotype and age groups (Supplementary Fig. 1A, B). Using qRT-PCR, we found significant upregulation of Gfap and Cd68 transcripts, well-established and commonly used markers of astrocytosis and microgliosis, in 9-month-old BPAN mice (Fig. 1G and H). We then looked at other pathology markers related to brain development or lysosomal/autophagy pathways, such as Lamp1, Mbp, Tfeb, Sort, Atg5, and Vamp4, but found no difference between BPAN Hemi mice and their littermate controls (Supplementary Fig. 1C).

Our experimental cohorts of female mice include Wdr45^+/+^ (WT), Wdr45^ex9+1g>a/+^ (Het) and Wdr45^ex9+1g>a/ex9+1g>a^ (Homo) mice (Supplementary Fig. 2). Unlike BPAN males, BPAN females did not show any differences compared to wild-type and heterozygous control littermates in the open field test at the age of 2 or 6 months (Supplementary Fig. 2B and C), in rotarod performance (Supplementary Fig. 2D) or weight (Supplementary Fig. 2E). We found no significant difference between genotype and age groups using glial markers Gfap and Cd68 via qPCR in brain tissue, except for elevation of microglial marker Cd68 in both Het and Homo mice at the age of 9 months (Supplementary Fig. 2F).

### CNS-targeted *WDR45* gene transfer rescued neurologic disease and brain pathology in symptomatic *WDR45* ^ex9+1g>a/Y^ mice

Most individuals with BPAN have already developed signs of neurologic disease, such as motor and cognitive deficits and seizures, before their diagnosis (5), and therefore gene therapy will have to be administered to the symptomatic patients. To address this in our preclinical proof-of-concept study, we set out to evaluate whether brain-targeted AAV-mediated *WDR45* gene transfer can rescue neurologic dysfunction in BPAN mice after symptom onset. To achieve broad CNS transduction and transgene expression in adult symptomatic mice, we designed an AAV-PHP.eB-WDR45 vector, since AAV-PHP.eB capsid results in broad brain transduction after systemic administration in mice (32). Human *WDR45* expression in this vector was driven by the synthetic promoter JeT, which is currently being used in a clinical trial for giant axonal neuropathy (GAN) (30, 33).

Only male mice were selected for this experiment since female homozygous mice showed no robust disease phenotype. AAV-PHP.eB-WDR45 (7e11 vg/mouse) was administered via tail-vein injection at 4 months of age, i.e, after the onset of hyperactive behavior in Hemi mice. 2 months later, the effect of the *WDR45* gene transfer was evaluated in the open field test (Fig. 2A), and no improvement of neurologic function was found in AAV-PHP.eB-treated BPAN mice compared to saline-treated littermates at this time (Fig. 2B). 5 months later, at 11 months of age, all the mice were re-tested in the open field, and correction of the hyperactive behavior was observed in the Hemi PHP.eB-WDR45-treated group compared to 6-month read-outs (Fig. 2B), showing the therapeutic effect of treatment at this later time point. No adverse events or weight changes were observed in the treated animals (Fig. 2C). qRT-PCR analysis of post-mortem brain tissue showed expression of human *WDR45* mRNA transcripts with the mean value of 1.25e5 copies/ug total RNA and 1.5 vector genome copies/cell (Fig. 2D). Western blot analysis showed that while levels of WIPI4 were lower than in healthy WT-saline samples, there was a significant increase in WIPI4 detection in the cerebral cortex of Hemi-treated vs. Hemi-saline control mice (Fig. 2E). Additionally, we have also observed correction of the Wipi2 levels in the cortical tissue of Hemi-treated vs. Hemi-saline control mice. In conclusion, these data provide proof-of-concept evidence that gene transfer of *WDR45* is efficacious in restoring neurologic and autophagy deficits when administered to symptomatic BPAN mice.

**Figure 2.**
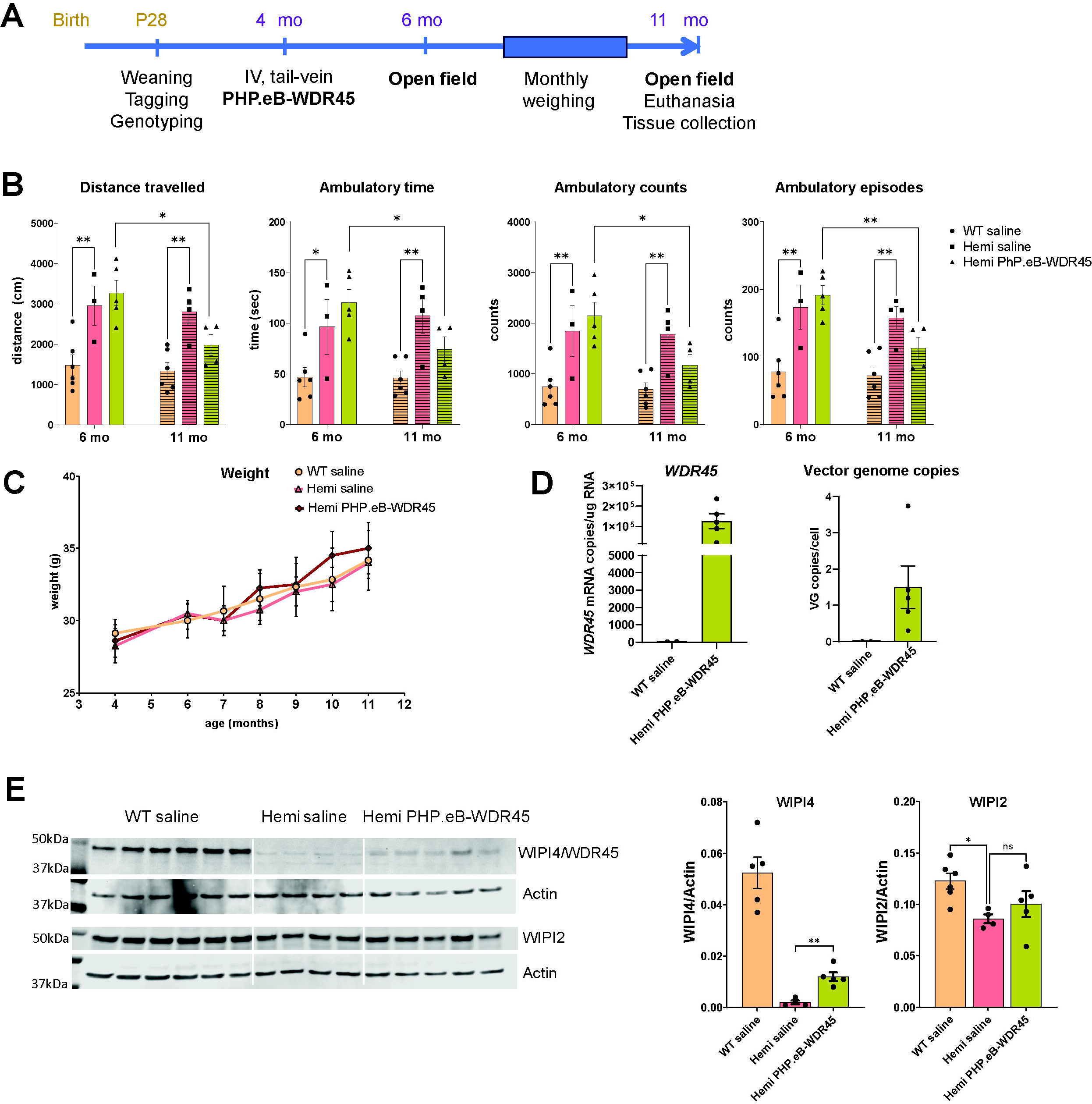
Intravenous administration of AAV-PHP.eB-WDR45 (PHP.eB-WDR45) in adult symptomatic Hemi BPAN mice corrected disease phenotype at a later stage of the disease. **A**. Schematic of experimental design. **B**. Open field test at 6 and 11 months of age shows a significant increase in activity, i.e., hyperactivity, in saline-treated hemi mice compared to WT-saline controls and significant correction of hyperactivity in Hemi mice treated with PHP.eB-WDR45 at 11 months. Data shown as individual values and average ± SEM; 6mo: n (WT-saline)=6, n (Hemi-saline) = 3 and n (Hemi-PHP.eB-WDR45) = 5; 11 mo: n (WT-saline)=6, n (Hemi-saline) = 4 and n (Hemi-PHP.eB-WDR45) = 4. Grouped statistical analysis was done using a two-way ANOVA (GraphPad Prizm) and Dunnett’s multiple comparisons test, **p<0.01. Comparisons of Hemi-PHP.eB values at 6 and 11 mo: * p<0.05, ** p<0.01 (Unpaired T-test). **C**. No changes in body weight have been observed in experimental groups. Data are shown as average values ± SEM; n (WT-saline) = 6-8, n (Hemi-Saline) = 4 and n (Hemi-PHP.eB-WDR45)=4-5. **D**. Quantification of the *WDR45* expression and vector genome copy number in the cerebral cortex of Hemi mice treated with PHP.eB-WDR45. Data are shown as average ± SEM, n (WT-saline) =2, n (Hemi- PHP.eB-WDR45)= 5. **E**. Western blot analysis of the brain tissue and quantification show increased levels of WIPI4 protein and Wipi2 in Hemi mice treated with PHP.eB-WDR45. Data shown as individual values and average ± SEM, *p<0.05, **p<0.01 (One-way ANOVA and multiple comparisons test).

### Neonatal intracerebroventricular (ICV) administration of scAAV9-WDR45 improves neurologic function and rescues brain pathology in *Wdr45* ^ex9+1g>a/Y^ mice

While AAV-PHP.eB shows high brain transduction with systemic administration, its ability to penetrate the blood-brain barrier is restricted to the species expressing the Ly6A receptor, with the highest transduction in C57Bl6 mice in which it was originally created and a very low brain transduction in non-human primates (NHP) (34, 35). Therefore, we next created a *WDR45* expression vector suitable for gene transfer in human patients based on the clinically tested self-complementary AAV9 vector (33, 36, 37). To drive the expression of WDR45, we used the same expression cassette as in our AAV-PHP.eB vector, i.e., with JeT promotor (30) and a synthetic short poly A signal (29). To achieve maximal vector transduction and broad transgene expression in the CNS tissue, we have performed intracerebroventricular (ICV) delivery of scAAV9-Jet-WDR45 vector (3e10 vg/mouse) to pups at post-natal day 1 (Fig. 3A). Heterozygous female and hemizygous male mice were used for breeding to obtain experimental cohorts that included wild-type (WT) and hemizygous (Hemi) males and heterozygous (Het) and homozygous (Homo) females. All litters were randomly assigned to either “vehicle” or “scAAV9-WDR45” treatment groups, and ICV injections were performed at P1 according to our established protocol. At weaning, mice were genotyped, same-sex mice were placed in the housing cages randomly within the treatment group, and only male mice were used for the follow-up evaluation. Neurologic function was evaluated in the open field test at 2 months of age. At 3 months of age, mice were sacrificed for tissue collection for post-mortem brain pathology and transgene expression analysis. ICV administration of scAAV9-WDR45 resulted in the expression of *WDR45* mRNA transcripts in the cerebral cortex of the treated Hemi mice with the mean value of 5.8e5 copies/ug total RNA and 1.5 vector genome copies/cell (Fig. 3B). ICV administration of scAAV9-WDR45 fully rescued hyperactivity behavior in the treated hemizygous mice (Fig. 3C). The treated mice have had no adverse effects or changes in body weight (Fig. 3D). Brain tissue analysis using Western blotting demonstrated a significant increase in WIPI4 detection (Fig. 3E). Our western blot data also showed significant recovery of the autophagy markers Wipi2 and LC3-II/I (Fig. 3E) further demonstrating efficacy of scAAV9-WDR45 in the BPAN brain tissue. Overall, our data demonstrates that brain-targeted AAV9-mediated gene transfer of *WDR45* rescues neurologic dysfunction and autophagy deficits in the BPAN mouse model.

**Figure 3.**
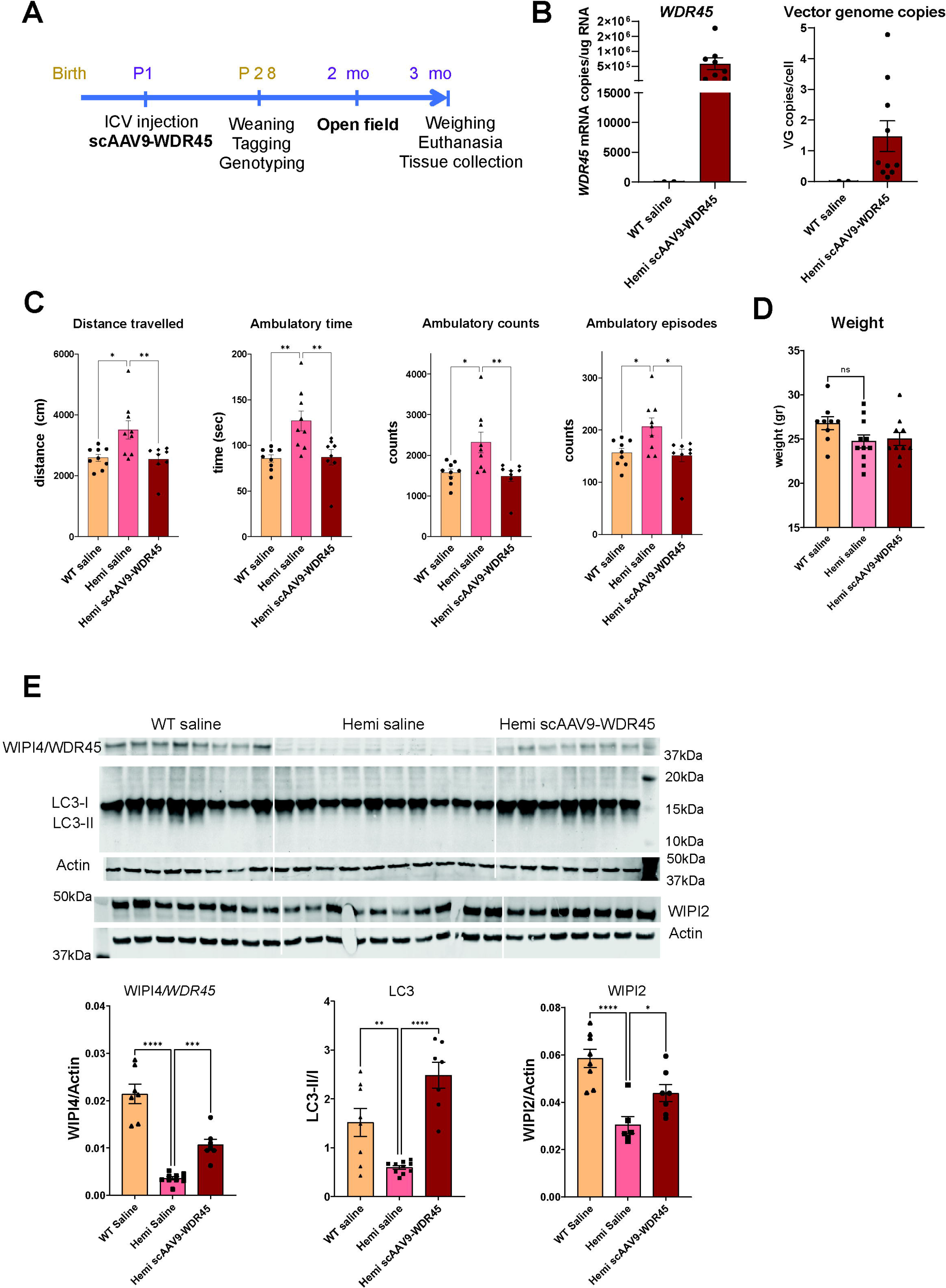
Neonatal ICV administration of scAAV9-WDR45 reduced hyperactivity and corrected autophagy markers in the brain tissue of hemizygous BPAN mice. A. Schematic of experimental design. B. Quantification of the *WDR45* expression and vector genome copy number in cerebral cortex of Hemi mice treated with scAAV9-WDR45. Data are shown as average ± SEM, n (WT-saline) =2, n (Hemi-scAAV9- WDR45)= 10. C. Open field test at 2 months of age shows a significant increase in activity, i.e., hyperactivity, in saline-treated hemi mice compared to WT-saline controls and significant correction of hyperactivity in Hemi mice treated with scAAV9-WDR45. Data shown as individual values and average ± SEM; n (WT-saline)=9, n (Hemi-saline) = 9 and n (Hemi-scAAV9-WDR45) = 8. *p<0.05, **p<0.01 (One-way ANOVA and multiple comparisons test). D. No changes in body weight have been observed in experimental groups at 3 months of age. Data shown as individual values and average ± SEM; n (WT- saline)=9, n (Hemi-saline) = 11 and n (Hemi-scAAV9-WDR45) = 10. One-way ANOVA and multiple comparisons test showed no difference between genotype/treatment groups. E. Western blot analysis of the brain tissue and quantification show increased levels of WIPI4, Lc3 and Wipi2 in Hemi mice treated with PHP.eB-WDR45. Data shown as individual values and average ± SEM, *p<0.05, **p<0.01, ***p<0.001, ****p<0.0001 (One-way ANOVA and multiple comparisons test), n (WT-saline)=8, n (Hemi- saline) = 10 and n (Hemi-scAAV9-WDR45) = 7. Three samples were excluded from Hemi-saline group in the Wipi2 data set dure to technical artefacts on the corresponding membrane.

### CNS-targeted overexpression of the lysosomal channel TRPML1 did not improve neurologic function in *WDR45* ^ex9+1g>a/Y^ mice

Since impairment of autophagy is thought to be a driving cause of pathogenesis in BPAN, we set out to test whether overexpression of the lysosomal channel TRPML1 can compensate for the loss of Wipi4 and ameliorate neurologic dysfunction in the mouse model of BPAN. TRPML1 is known as a master regulator of lysosome-related pathways and processes (reviewed in (38)), including autophagy induction through TFEB and mTORC1 (39, 40), is also involved in autophagophore formation via induction of the Beclin1/VPS34 autophagic complex and the generation of phosphatidylinositol 3-phosphate (PI3P) (41). Previously, we have developed AAV vectors expressing human TRPML1 (encoded by MCOLN1) and successfully demonstrated preclinical efficacy in the mouse model of Mucolipidosis type IV, a deficiency of TRPML1 (42). Here, we used one of these vectors, scAAV9-MCOLN1, to test whether overexpression of TRPML1 in the mouse model of BPAN will be sufficient to correct disease phenotype. To minimize the number of animals and allow for direct comparisons, this experiment was conducted in parallel with testing scAAV9-WDR45 via ICV injections in neonatal BPAN mice (Fig. 3). Additional litters were dosed with scAAV9-MCOLN1 at 2e10 vg/mouse, a dose that resulted in full rescue of the phenotype in the MLIV mouse model in our previous study (42). Unfortunately, despite the high expression of MCOLN1 transcripts in the cortical tissue of BPAN mice, scAAV9-MCOLN1 failed to rescue hyperactivity (Fig. 4B, C). Based on this data, we conclude that despite the broad functional impact of TRPML1 in regulating autophagy, its solo targeting is unlikely to be a promising therapeutic strategy for BPAN.

**Figure 4.**
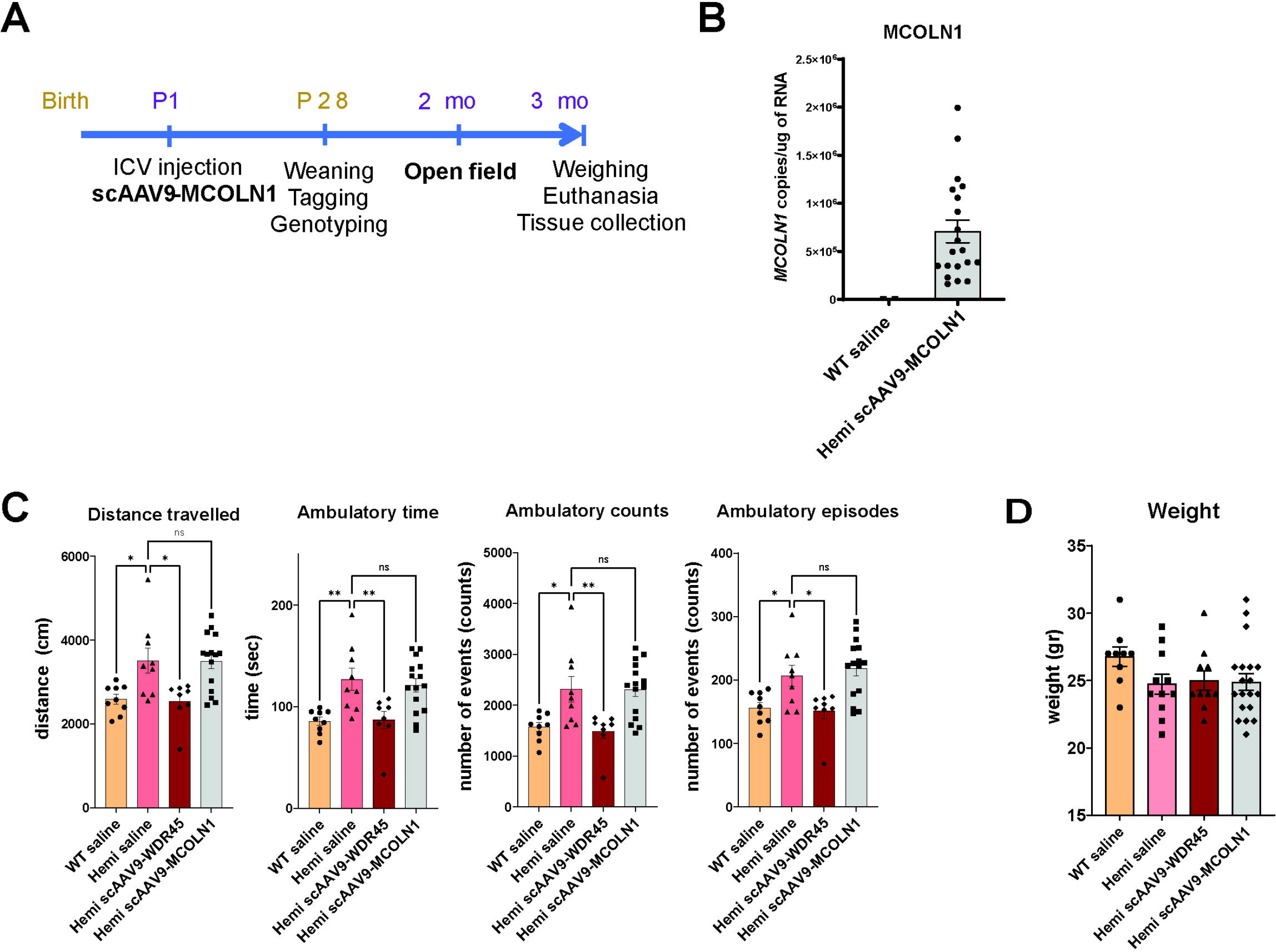
Neonatal ICV administration of scAAV9-MCOLN1 failed to correct disease phenotype in hemizygous BPAN mice. A. Schematic of experimental design. B. Quantification of the MCOLN1 expression in cerebral cortex of Hemi mice treated with scAAV9-MCOLN1. Data are shown as average ± SEM, n (WT-saline) =2, n (Hemi-scAAV9-WDR45) = 20. C. Open field test at 2 months of age shows a significant increase in activity, i.e., hyperactivity, in saline-treated hemi mice compared to WT-saline controls, significant correction of hyperactivity in Hemi mice treated with scAAV9-WDR45, and lack of correction in Hemi mice treated with AAV9-MCOLN1. Data shown as individual values and average ± SEM; n (WT-saline)=9, n (Hemi-saline) = 9 and n (Hemi-scAAV9-WDR45) = 8, n (Hemi-scAAV9- MCOLN1)=15. *p<0.05, **p<0.01 (One-way ANOVA and multiple comparisons test). D. No changes in body weight have been observed in experimental groups at 3 months of age. Data shown as individual values and average ± SEM; n (WT-saline)=9, n (Hemi-saline) = 11 and n (Hemi-scAAV9-WDR45) = 10, n (Hemi-scAAV9-MCOLN1)=19. One-way ANOVA and multiple comparisons test showed no difference between genotype/treatment groups.

## Discussion

BPAN is an ultra-rare severe disease with early neurologic and cognitive presentation in the form of delayed motor development, seizures, and autism-like symptoms and a progressive course leading to a movement disorder and dementia in the second and third decades of life. It was first described in 2012 (1). With a growing community of patients worldwide, the unmet medical need in BPAN is vast, with no therapies or interventional clinical trials registered at www.Clinicaltrials.gov. While the pathological cascade in this disease is mainly associated with disruption of autophagy due to loss of function of the autophagy scaffold protein WIPI4, the precise pathophysiological mechanisms are currently unknown. Several mouse models have been developed and characterized to better understand the role of Wipi4 in brain development and function. They include a constitutive *WDR45* knock-out model with a stop codon in exon 6 (23), a constitutive knock-out model generated by crossing *WDR45* fl/Y mice with Zp3-Cre line (24), and a constitutive *WDR45* knock-out TALEN mouse (25). Conditional models include a pan-neuronal *WDR45* knock-out model (26) and a dopaminergic neuron-specific WDR45^Fl/Fl^/DAT^CreERT2^ knock-out model (27). These models showed progressive neuropathologic features in the brain, including microgliosis and astrocytosis in some brain regions, axonal swelling and degeneration, loss of cerebellar Purkinje cells, defective autophagy, and accumulation of p62/SQSTR aggregates with various ages of onset in different models and the earliest signs reported at 4 months. Cognitive deficits (impaired spatial and contextual memory), autism-spectrum disorders-like phenotype, and locomotor deficits have been detected in BPAN mouse models at different ages: in full-body *WDR45* knock-out mice by Wan et al (23), both motor and cognitive deficits were reported at 6-8 months, with the exception of the reduced marble burying behavior and a higher sensitivity to pilocarpine-induced seizures observed in KO mice at 2 months; in the TALEN *WDR45* KO model, statistically significant motor and cognitive deficits were observed after 11 months of age (23, 25). Both CNS-specific *WDR45* knock-out models, Nes-Wdr45^Fl/Y^ and Wdr45^Fl/Fl^/DAT^CreERT2^ mice, demonstrated the onset of disease phenotype after 11 and 17 months, correspondingly (26, 27), suggesting milder phenotype in the conditional models. In addition to the neurodegenerative phenotypes described earlier, Wipi4 was recently shown to play a role in brain development (43). In mouse brain development, expression of Wipi4 was shown to be the highest at embryonic development, dropping after post-natal day 8 and reaching the lowest level by 3 months of age. Acute knock-down of *WDR45* led to abnormal dendritic development and synaptogenesis supporting its role in brain development and further linking loss of WIPI4 function with neurodevelopmental abnormalities seen in individuals with BPAN. Overall, despite the enthusiastic effort generating and characterizing these mouse models, late or subtle phenotypes precluded them from being strong disease-relevant models of BPAN that can be used for preclinical testing of drug candidates.

In this study we have characterized a novel mouse model of BPAN, *WDR45* ^ex9+1g>a^ mouse. *WDR45* ^ex9+1g>a^ mouse mimics a human pathogenic variant c.830 + 1G>A reported in the girl that, due to clinical presentation, was initially diagnosed with Rett syndrome with childhood iron deposition in brain (44). *WDR45* ^ex9+1g>a/Y^ mice demonstrate early and significant hyperactivity behavior corresponding to attention deficit and hyper-arousal in pediatric patients with BPAN. Unlike in other previously characterized models of BPAN, the behavioral changes in *WDR45* ^ex9+1g>a/Y^ mice in the form of increased traveled distance, ambulatory time, ambulatory episodes, and ambulatory counts, all measured in the open field test were detected early, starting at 2 months of age, and were persistent through the course of the disease as evident by the presence of this phenotype at 11 months of age. To our knowledge, this is the first report of the early phenotype in a BPAN mouse model directly relevant to human disease. The observed hyperactivity readouts provide valuable disease-relevant outcome measures for preclinical therapeutics testing in translational and therapy-oriented studies of BPAN.

On the molecular level, we found that *WDR45* ex9+1g>a variant led to reduced levels of Wipi2 protein in the brain homogenates from hemizygous mice as compared to their wild-type littermates. WIPI2 plays an essential and conserved role in autophagosome formation and functions in tandem with WIPI1 in the early stages of autophagosome formation upstream of WIPI4 (45). WIPI2 binds PI3P and enables LC3- lipidation and autophagophore formation via recruitment of ATG12-5-16L1 complex (45), and its knockdown leads to near complete inhibition of the autophagic flux (18, 20). WIPI4, in response to AMPK stimulation, binds ATG2 and facilitates the elongation of the autophagosomal membrane (18). Unlike WIPI2, the knockdown of WIPI4 alone leads to only a slight reduction of the autophagy, affecting mostly the size of the autophagosomes (20). The interaction of WIPI4 and WIPI2 was reported (18), and a recent study demonstrated the essential role of WIPI2 in supporting the function of WIPI4 in autophagy (20). Given the role of WIPI2 in autophagy and its interactions with WIPI4, we assessed how loss of Wipi4 affected Wipi2 levels in the brain tissue in *WDR45* ^ex9+1g>a/Y^ mice. We found a slight but significant reduction of Wipi2 levels in brain homogenates from *WDR45* ^ex9+1g>a/Y^ mice compared to their wild-type counterparts at all time points. To our knowledge, this is the first evidence of reduced Wipi2 in BPAN animal model, as Wipi2 levels were not reported in other published models of BPAN (23, 24, 25, 26, 27). Importantly, we also showed that gene transfer of *WDR45* led to partial but significant correction of the Wipi2 levels in the brain tissue. Given the established functional interactions of WIPI4 and WIPI2 in autophagosome formation, the subtle effect of WIPI4 loss on autophagy in cell and animal models, and the low sensitivity of the canonical autophagy assays, our findings suggest that WIPI2 can be used as a molecular surrogate marker to assess WIPI4 loss of function and its recovery in the mouse model of BPAN.

In this study using the *WDR45* ex9+1g>a/Y model of BPAN, we set out to evaluate whether the pathological manifestation of BPAN can be reversed or attenuated by AAV-mediated gene transfer of human WDR45. We started our investigation by testing whether brain-targeted expression of *WDR45* would have a therapeutic effect in the symptomatic adult mice which would mimic in the animal model the clinical settings for a future first-in-human trial in BPAN patients, all of whom will be recruited after symptom onset. To achieve brain-targeted expression of WDR45, we employed an AAV-PHP.eB capsid. It was developed using in vivo-directed evolution of the AAV9 capsid by the Cre REcombinase-based Targeted Evolution (CREATE) system capsid and was shown to have higher brain transduction and in particular, neurons over its parental capsid AAV-PHP.B in adult C57Bl6 mice after systemic administration, resulting in transduction of 69% of cortical and 55% of striatal cell with 1e11 vg/animal (46). To drive the expression of WDR45, we have selected a short synthetic promotor JeT that leads to ubiquitous and mild expression of the *WDR45* transgene upon AAV-mediated delivery into cells (30). This design aimed at delivering *WDR45* broadly to the brain tissue but maintaining its expression at the modest – physiological or supraphysiological – levels to avoid overexpression-induced toxicity and cellular stress, especially given that WIPIs are not secreted and no cross-correction between transduced and non-transduced brain cells was expected.

Administration of AAV-PHP.eB-WDR45 in adult *WDR45* ^ex9+1g>a/Y^ mice at 4 months of age led to a reduction of hyperactivity at 11 months of age, suggesting correction of disease phenotype at the late stages of the disease (Fig. 2). This is an important outcome, which implies that delivery and expression of the functional *WDR45* transcript can alleviate neurologic symptoms even if the intervention took place after symptom onset. Despite this promising outcome in 11 months-old mice, open field test at 6 months of age, i.e. 2 months post-treatment, showed no therapeutic effect in hyperactivity phenotype in *WDR45* ^ex9+1g>a/Y^ - PHP.eB-WDR45 treated animals compared to saline-treated Wdr45^ex9+1g>a/Y^ group. Together with the Western blot data showing only a modest increase in WIPI4 expression in the brain tissue (which was 4.3-fold lower than endogenous expression of Wipi4 in the brain of saline-treated wild-type mice), this slow correction might be explained by lower than optimal protein expression levels resulting in a slow restoration of WIPI4-dependent pathways and neurologic function.

Despite the great BBB penetrance and CNS-targeting properties of the AAV-PHP.eB in mice, they do not translate to other species, limiting the clinical translatability of this vector (47). Therefore, in the next set of experiments, we designed and tested a self-complimentary AAV9-WDR45 expressing vector using the same expression cassette as AAV-PHP.eB-WDR45. scAAV9 is currently the vector of choice for CNS indications due to a higher tropism for neurons than in any other naturally occurring AAV capsids (48, 49, 50). However, to achieve broad brain biodistribution and minimize the risk of high-dose toxicity with intravenous delivery, intra-CSF administration is required (51). While different routes of intra-CSF delivery of AAVs, i.e. intrathecal lumbar puncture, intra-cisterna magna or intracerebroventricular (ICV) in the lateral ventricles, can be used in the large animal models and in the clinical setting, in the mouse neonatal ICV injections serve as a surrogate intra-CSF delivery method to achieve the broad brain biodistribution and intra-neuronal delivery (33, 52, 53), and we have used this route of delivery in our study to deliver scAAV9-WDR45 into the CNS of *WDR45* ^ex9+1g>a/Y^ mice. Importantly, this early post-natal intervention resulted in full prevention of the hyperactivity phenotype in scAAV9-WDR45-treated *WDR45* ^ex9+1g>a/Y^ mice at two months of age (Fig. 3), providing proof-of-concept evidence that intervention with a clinically validated AAV vector in the mouse model of BPAN leads to the therapeutic improvement of neurologic function. In addition to functional rescue, we have also observed a partial but significant increase in the levels of Wipi2, and correction of the autophagy marker Lc3-II indicative of improvement of autophagy in the scAAV9-WDR45-treated *WDR45* ^ex9+1g>a/Y^ mice. Based on Western blot analysis, expression of WIPI4 in the brain tissue of scAAV9-WDR45-treated *WDR45* ^ex9+1g>a/Y^ was 3-fold higher than expression of the remaining detectable Wipi4 protein in the saline-treated *WDR45* ^ex9+1g>a/Y^ mice but remained at supraphysiological levels and was 2-fold lower than endogenous Wipi4 expression in the saline-treated wild-type brain tissue. Our data measuring *WDR45* mRNA transcript, vector genome copies, Wipi4 protein levels in correspondence with the administered vector dose (3e10vg/mouse), and efficacy outcomes in the mouse model of BPAN set the stage for the future dose-optimization and safety translational study for BPAN. Overall, these data indicate that AAV-mediated gene supplementation is a promising therapeutic strategy for BPAN, a disease with extremely high unmet medical need for which no therapies currently exist.

In parallel with *WDR45* gene transfer proof-of-concept study in *WDR45* ^ex9+1g>a^ BPAN mouse model, we have tested whether AAV-mediated CNS targeted overexpression of the lysosomal channel transient receptor potential mucolipin-1, or TRPML1 (encoded by MCOLN1 gene) is capable of alleviating disease phenotype in BPAN mice. Studies of the lysosomal channel TRPML1 revealed its important role in regulating many aspects of lysosomal biology. Most of the known functions of TRPML1 are related to its role in Ca^2+^ transport and include the fusion of membrane organelles such as endosomes and autophagosomes with the lysosome, fusion of lysosomes with the plasma membrane for lysosomal exocytosis and phagocytosis, and Ca^2+^-driven regulation of the transcription factor EB (TFEB), a master regulator of the lysosomal gene network (38, 54, 55, 56, 57, 58, 59, 60). In addition, a direct role of TRPML1 in regulating the activity of the key metabolic signaling regulator, mTOR (60, 61, 62), and a role in autophagophore formation through a TFEB-independent mechanism (41) have also been recently documented, further highlighting the importance of TRPML1 in autophagy regulation. Loss of TRPML1 function in humans results in mucolipidosis IV (MLIV), an autosomal-recessive disorder with severe early CNS disability, vision loss, achlorhydria, i.e., reduced secretion of chloric acid in the stomach, and brain iron accumulation in the basal ganglia (OMIM #252650) (63). Given a broad array of TRPML1 functions to regulate lysosome-related pathways and processes, its recently discovered role in autophagophore formation via induction of the Beclin1/VPS34 autophagic complex and the generation of phosphatidylinositol 3-phosphate (PI3P) (41), coupled with its well-established role in autophagy induction through TFEB (39, 40), we hypothesized that upregulating TRPML1 expression could compensate for deficient autophagy resulting from loss of WIPI4 and in rescue of the disease phenotype in the BPAN model. To test this, we have used scAAV-MCOLN1 vectors previously shown to rescue disease phenotype in the MLIV mice (42) via neonatal administration into *WDR45* ^ex9+1g>a/Y^ mice. Although neonatal ICV administration of scAAV9-MCOLN1 resulted in brain biodistribution and expression of the transgene, we have observed no rescue of the hyperactivity behavior in the scAAV9-MCOLN1-treated *WDR45* ^ex9+1g>a/Y^ mice as compared to their saline-treated counterparts, suggesting that overexpression of MCOLN1 might not be sufficient to see the therapeutic rescue in BPAN. Based on our data, CNS-targeted AAV-mediated gene transfer of *WDR45* is a promising therapeutic strategy for BPAN.

## Conclusions

In conclusion, we performed characterization of the novel *WDR45* ^ex9+1g>a^ mouse model for BPAN and identified hyperactivity as a disease-relevant functional phenotype that can be used for preclinical assessment of therapeutic interventions. On the molecular level, we found a reduced abundance of the two autophagic markers in the brain tissue of BPAN mice, Lc3 and Wipi2, indicative of inhibition of autophagosome formation. Importantly, unlike previously published models of BPAN, these phenotypical characteristics developed in the BPAN mouse model early in the course of the disease. Given the early onset of BPAN disease in human patients, this model is advantageous for preclinical trial design and outcome interpretation in translational and therapy-focused studies. We used this mouse model for a proof-of-concept preclinical study to evaluate whether AAV-mediated CNS-targeted gene transfer of *WDR45* can provide therapeutic benefit and be considered a therapeutic paradigm for BPAN. We observed successful expression of human *WDR45* transcripts and WIPI4 protein in the brain tissue, rescue of hyperactive behavior, and correction of autophagy markers in the brain tissue. This data demonstrates that *WDR45* gene transfer can be a promising therapeutic strategy for BPAN if efficacious expression levels can be achieved in the brain.

## Acknowledgments

The authors are grateful to Christina Mascarenhas, Kelly Kozole (Don’t Forget Morgan Foundation), and Bill Roth (Harper’s Hope Foundation) for their efforts to promote research and find a cure for BPAN; and provide financial support for this study. The authors are also grateful to Dr. Kristin Grimsrud for early discussions and preliminary assessment of *WDR45* ex9+1g>a mouse phenotype performed in the UC Davis Department of Pathology and Laboratory Medicine.

## Authorship confirmation/contribution statement

M.C.C.- Formal Analysis, Methodology, Investigation, Visualization, Writing – original draft; C.E.S.- Investigation, Visualization, Writing – review & editing; M.M.B.- Investigation, Visualization, Writing – review & editing; M.L.S.- Investigation, Methodology, Validation, Writing – review & editing; U.C.- Methodology, Formal Analysis, Writing – review & editing; B.M.- Resources, Writing – review & editing; JL.P. - Resources, Writing – review & editing; Y.G.- Conceptualization, Funding acquisition, Methodology, Project administration, Supervision, Visualization, Writing – original draft, Writing – review & editing.

## Author(s’) disclosure

Authors disclose no conflict of interest.

## Funding statement

This work was funded by research grants from Don’t Forget Morgan and Harper’s Hope to Y.G.

## Notes

### Competing Interest Statement

This work was funded by research grants from Dont Forget Morgan and Harpers Hope to Y.G.

