## Supplementary Info for "AAV-mediated gene transfer of WDR45 corrects neurologic deficits in the mouse model of beta-propeller protein-associated neurodegeneration"

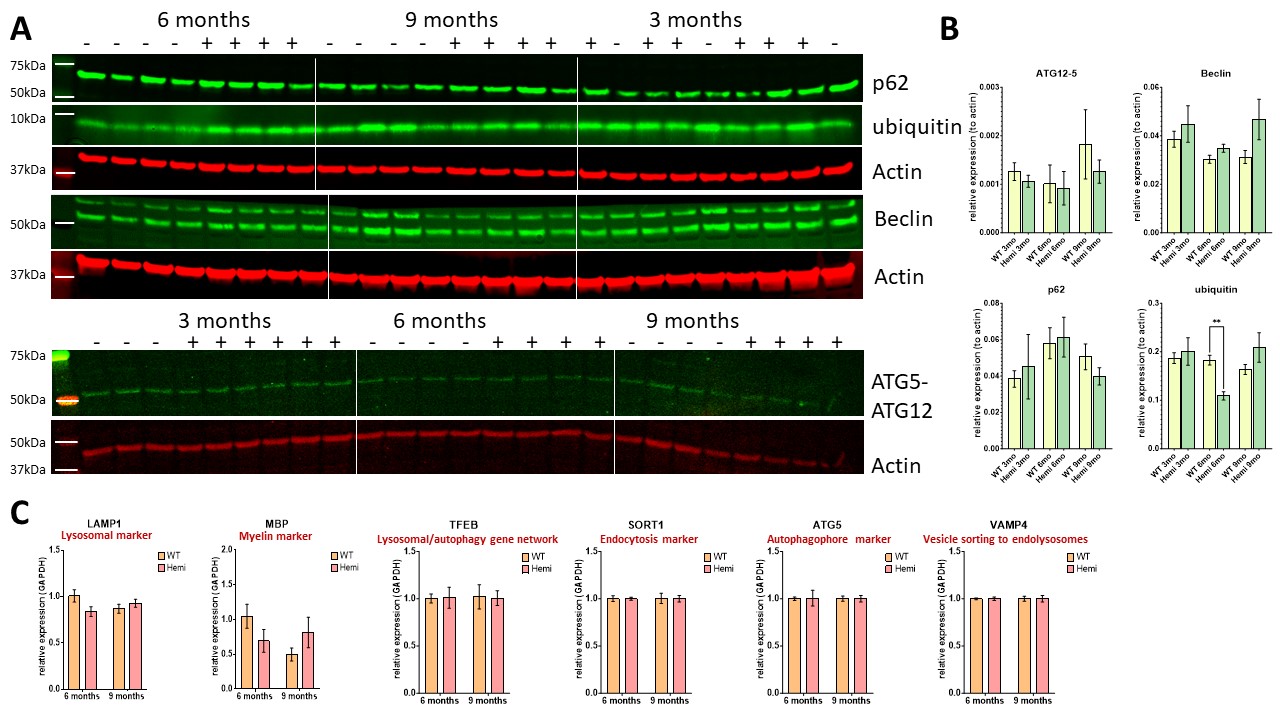

**Supplementary Figure 1.** Western blot (A) and quantification (B) of autophagy and protein degradation markers in cerebral cortex of Hemi (-) and WT (+) mice at 3, 6 and 9 months of age. Data are shown as average ± SEM; n=6 (WT) and 3 (Hemi) at 3 mo, n= 4 per genotype at 6 and 9 mo. C. qRT-PCR analysis of a panel of markers in Hemi and WT cortex at 6 and 9 months. Data are shown as average ± SEM, n= 4 per genotype and age group.

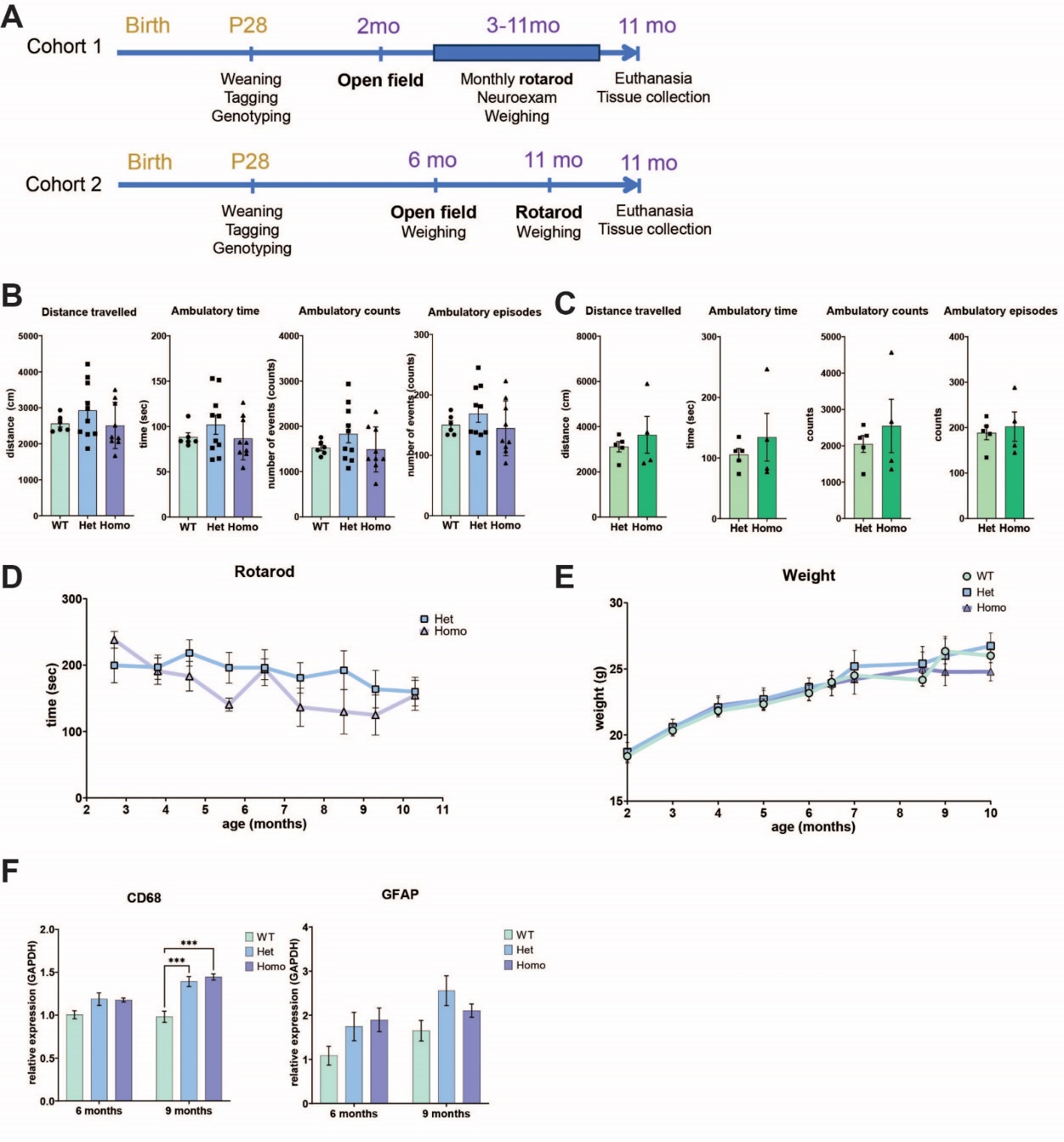

**Supplementary Figure 2.** Characterization of disease phenotype in *Wdr45^ex9+1g>a^* female mice. **A**. Schematic representation of the experimental design. B, C. Open field test shows no significant difference between *Wdr45^ex9+1g>a/ ex9+1g>a^* (Homo)*, Wdr45^ex9+1g>a/+^* (Het) and *Wdr45 ^+/+^* (WT) mice at either 2 (**B**) or 6 (**C**) months of age using ordinary one-way ANOWA and Unpaired T-test, correspondingly. 2mo: n (Homo)=9*,* n (Het) = 10 and n (WT) = 6; 6 mo: n (Homo) = 4, n (Het) = 5. Data shown as individual values and average ± SEM. **D**. Monthly rotarod testing shows no significant changes between Het and Homo female mice from 3 to 11 months of age. Data are shown as average values ± SEM; n (Het) = 7 (3-10 mo) and 12 (11 mo), n (Homo) = 5 (3-10 mo) and 10 (11 mo). Grouped statistical analysis using a mixed-effects model with the Geisser-Greenhouse test (GraphPad Prizm): p (age)= 0.041; p (genotype) = 0.47. Sidak’s multiple comparisons test showed no significant differences between WT and Hemi at any time point. **E**. No significant changes in body mass have been observed between Homo (n=9-14), Het (n=7-16) and WT (n=5-6) mice from 2 to 10 months of age. Data are shown as average values ± SEM; Grouped statistical analysis using a mixed-effects model with the Geisser-Greenhouse test (GraphPad Prizm): p (age) <0.0001; p (genotype) = 0.62. Sidak’s multiple comparisons test showed no significant differences between WT and Hemi at any time point. **F**. qRT-PCR analysis of the activated microglia/macrophage marker CD68 and astrocyte marker GFAP in cerebral cortex at 9 months. Data are shown as average ± SEM, ***p<0.001; two-way ANOVA (p (age) = 0.0028, p (genotype) <0.0001) and Tukey’s multiple comparisons test ; n= 4 per genotype and age group.
